# A Meta-Analysis of Executive Functions in Frontal Cortex: Comparing Healthy and Neuropsychiatric Groups

**DOI:** 10.1101/335109

**Authors:** Abigail B. Waters, Lance P. Swenson, David A. Gansler

## Abstract

**Background:** The neural architecture of executive functions remains a topic of considerable clinical and academic interest in the clinical neurosciences, given its strength as a transdiagnostic predictor of adaptive functioning with high heritability. In recent years, meta-analyses have shown a consistent relationship between prefrontal cortex size and executive functioning task performance in healthy adults and lesion patients, with increases in measures of cortical size (i.e., volume or thickness) associated with better executive functioning performance. There is a gap in meta-analytic literature assessing these relationships in neuropsychiatric populations, their effects relative to healthy controls, and differential contributions of brains regions and neuropsychological paradigms.

**Methods:** We conducted a meta-analysis of published studies (*k* =30) that assessed the relationship between executive functions and frontal regions *in vivo* (*N* = 1935) for both healthy (20 samples) and neuropsychiatric (21 samples) adults. Random effects modeling was used to calculate mean effect sizes and CIs.

**Results:** Larger volumes and thickness were associated with better executive functioning in both healthy (r =.35, 95% CI =.29 -.39) and neuropsychiatric populations (r =.47, 95% CI =.40 -.51), with the effect size for neuropsychiatric populations being significantly larger compared to healthy controls. While there was variability between tasks, there were no significant differences in effect size between neuropsychological paradigms or brain region classification.

**Conclusions:** These results indicate the relationship between healthy adult performance on neuropsychological testing is less associated with cortical size compared to neuropsychiatric adults.

## Introduction

The neural architecture of executive functions (EF) remains a topic of considerable clinical and academic interest in the clinical neurosciences. Poor EF, as measured by neuropsychological testing, has been associated with lowered instrumental activities of daily living(1), general cognitive decline(2), and increased mortality(3), validating its strength as a transdiagnostic predictor of clinical outcomes of interest. These tests have been historically labelled “frontal tests” as validated by lesion studies(4) and have been correlated with measures of cortical size and health in the frontal lobe(5, 6). These EF are influenced by a highly heritable general factor, not related to intelligence or processing speed(7), and as the effect of genes on cognitive phenotype is not directly observable, the cerebral cortex is a considerable mediator of that relationship. However, given high variability in constructural understanding of EF and sample populations, meta-analytic review of findings are necessary to consolidate and interpret the range of results.

There is considerable debate about the functional and cortical organization of EF. EF are broadly characterized as the set of cognitive processes needed to regulate, coordinate, and plan behavior(8). In previous research, the construct has been variously defined as a set of distinct, separate skills(9), a more general unitary factor(10), or reflecting both unitary and diverse influences(11). There is support in the literature for both of these positions from factor analytic and imaging studies(12, 13). The debate regarding the functional organization of EF parallels that on functional heterogeneity of the prefrontal cortex(14). However, it is generally agreed upon that these adaptive functions are often used in conjunction with one another and exist as part of a larger system of cognitive control.

In both healthy and patient populations, EF is commonly assessed with neuropsychological paradigms, such as the Wisconsin Card Sort (WCST), Stroop Color Word Interference Test (CWI), the Trail Making Test (TMT), Verbal Fluency (VF), and working memory (i.e., span) tasks (WM). There has been consistent evidence that EF tasks are sensitive to the presence of brain damage, but the degree to which these abilities can be localized to the prefrontal cortex in the context of lesion studies is less clear(15). Although early lesion studies emphasized focalization of EF tasks to dorsolateral, orbitofrontal, and ventromedial regions in the frontal lobe, more contemporary reviews of lesion studies have not supported a one-to-one relationship between any task and region(4). Integrating previous findings that have shown the WCST associated with dorsolateral damage specifically(16), and the TMT with inferior medial regions(17), another interpretation emphasizes that executive deficits observed in patients are the result of multiple attentional circuits connecting both frontal and posterior regions, which are adaptive to multiple contexts(15).

Although lesion studies have been instrumental in mapping the functional organization of EF in patient populations, investigations of these relationships in non-clinical populations are important for understanding normal neurocognitive functions as a baseline of comparison. While often studies compare patients with healthy controls in terms of performance, it is imperative to examine whether the degree to which performance is influenced by brain size varies by group.Like in clinical populations, tasks of EF have been linked with specific subregions of prefrontal cortex in healthy adults. However, meta-analysis in healthy populations shows that while some tasks are more associated with volume than others, there is little evidence to support focalized organization of the prefrontal cortex for executive functions as measured by clinical neuropsychological tasks(14). A 2014 meta-analysis(6), examined the relationship between prefrontal cortex volume and thickness with EF in healthy adults. Larger volume and cortical thickness were associated with better performance on tasks of EF. However, significant brain-behavior associations were only found in the lateral and medial, and not orbital, regions of the prefrontal cortex. It is possible that these variable findings result from smaller contributions of brain size for healthy populations, relative to protective factors like education or cognitive reserve. Examining these relationships in the context of both community dwelling and neuropsychiatric populations via meta-analysis can further clarify how these tasks and regions are linked, and the comparative contribution of cortical size for tasks of EF for these two groups.

*Research Question 1.* Is there a significant positive association between cortical volume and thickness and performance on tasks of EF?

*Research Question 2.* Does the magnitude of that association vary as a function of task or brain region distribution?

*Research Question 3.* Is there a difference in the magnitude of effect sizes between healthy and neuropsychiatric groups?

*Research Question 4.* Are there differences in the magnitude of effect sizes between neuropsychiatric groups?

## Methods and Materials

### Selection of Studies

A literature search of the computerized database PubMed was conducted in January 2017 using the following key words: (frontal OR prefrontal) AND (volume OR volumetric OR atrophy OR cortical thickness OR cortical thinning OR morphometry OR FreeSurfer) AND (Executive OR WCST OR Wisconsin Card OR Stroop OR Trail Making Test OR TMT OR Verbal Fluency OR Working Memory OR Iowa Gambling Task OR IGT).

### Inclusion Criteria

The search items described above produced a list of 1,404 research items (see Figure 1). Titles and abstracts were examined to determine their appropriateness for the current study, and 248 studies were retained for further consideration. Case studies, research of non-human subjects, and human participants under age 18 were excluded.

**Figure 1.**
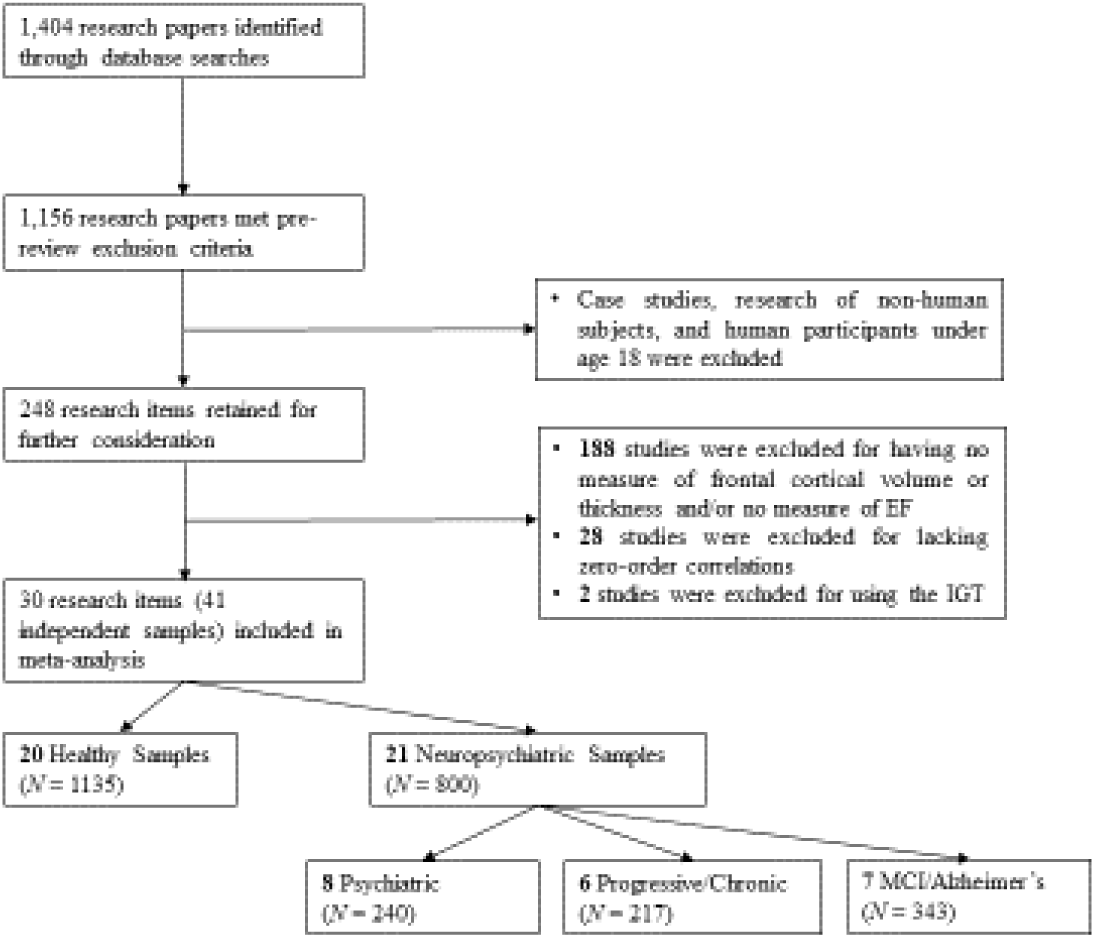
Flow chart describing identification and screening of articles and studies.

Studies were selected if they measured frontal cortical volume or thickness, included at least one measure of EF, and contained usable statistics (e.g., correlation) relating variables of interest in human adults. 30 studies were retained in the final selection of meta-analyses. Of the 30 studies, 3 examined cortical thickness and 27 examined cortical volume (Table 1). Cortical thickness and volume were collapsed into a single indicator for analyses, as previous meta-analysis has shown their effect sizes are equivalent(6). Multiple samples were pulled from 9 studies; however, all 41 effect sizes represent independent observations. The final sample (*N* = 1935) included data for both healthy (20 samples) and neuropsychiatric (21 samples) adults.

**Table 1.**
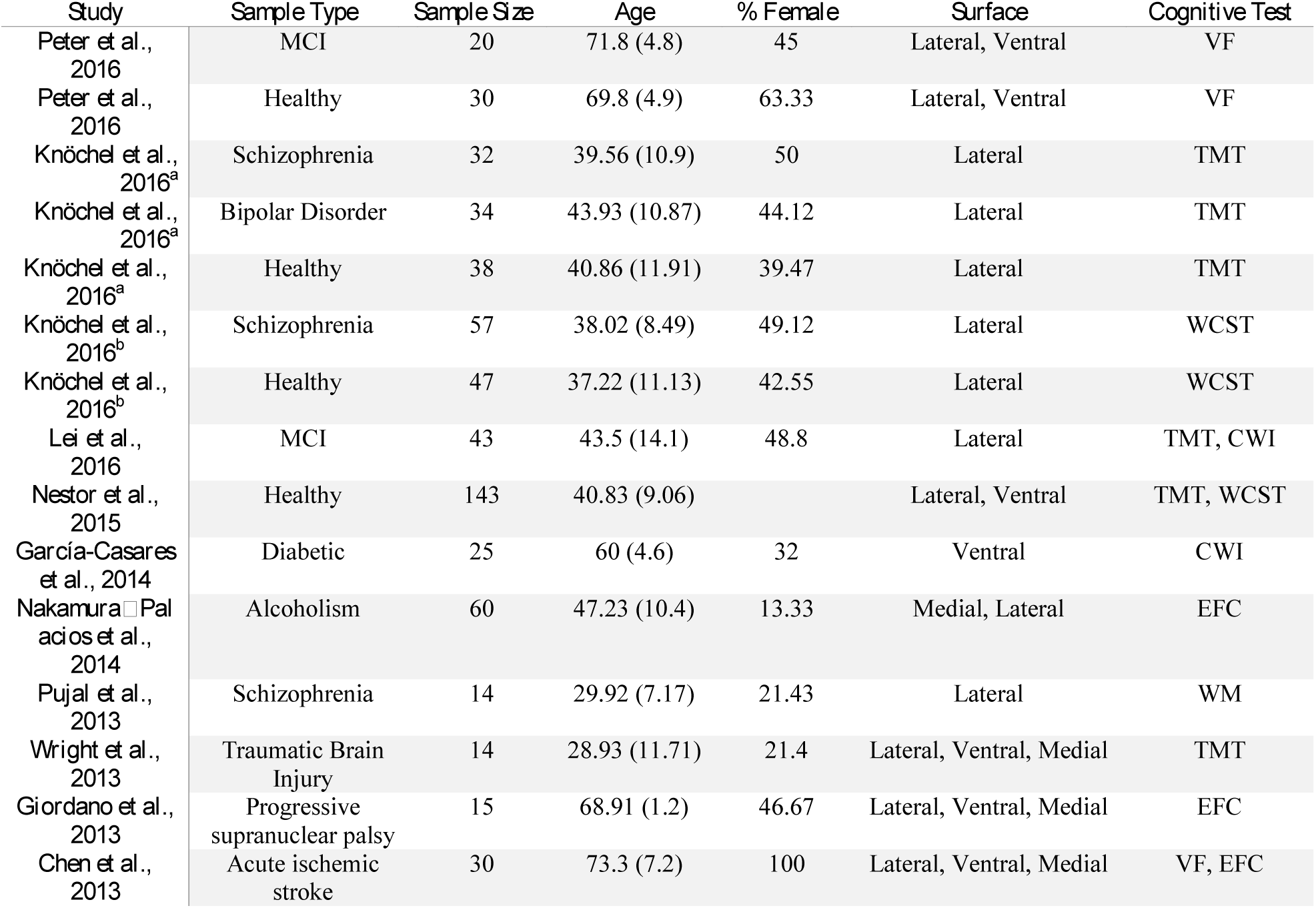
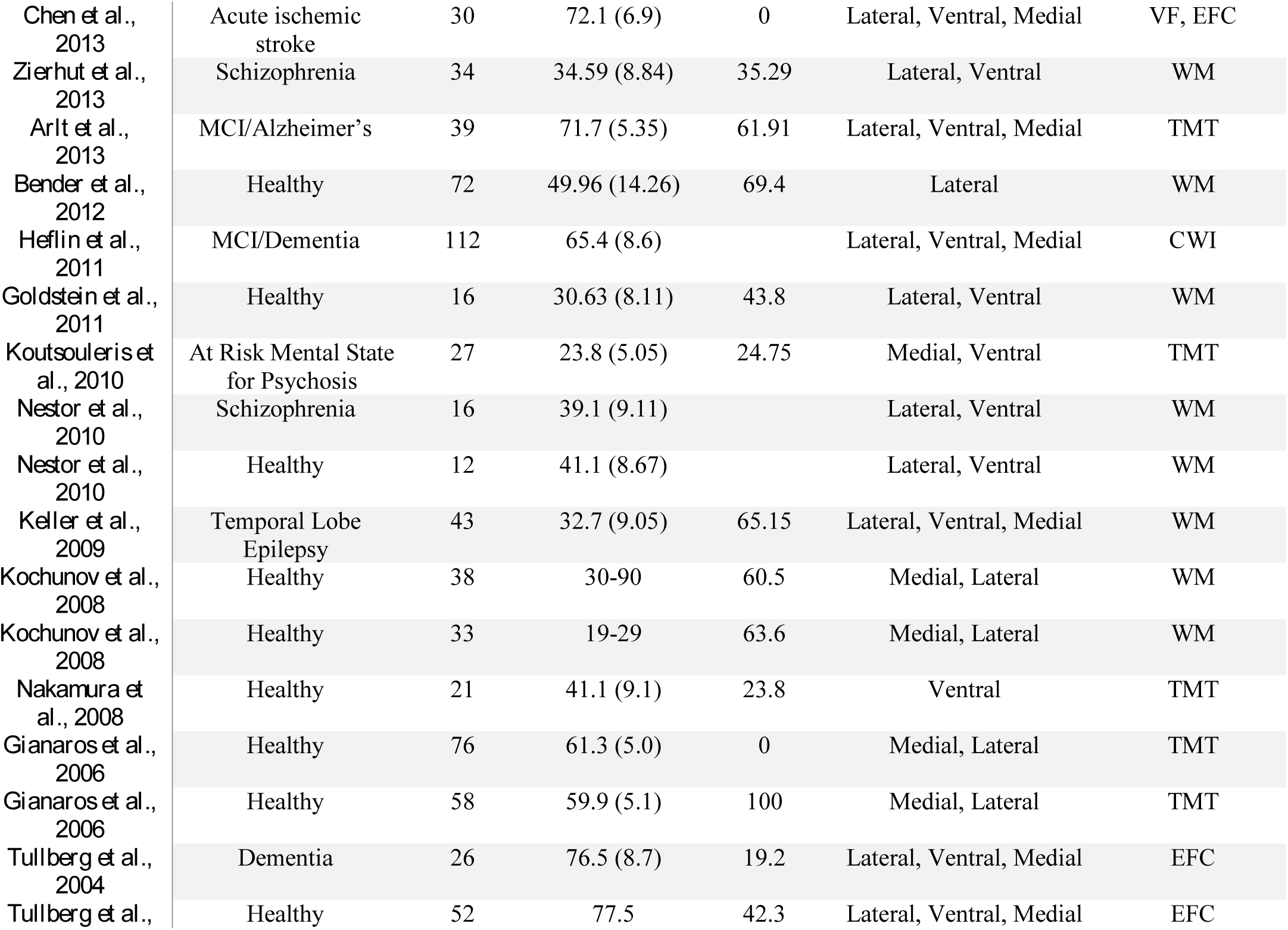
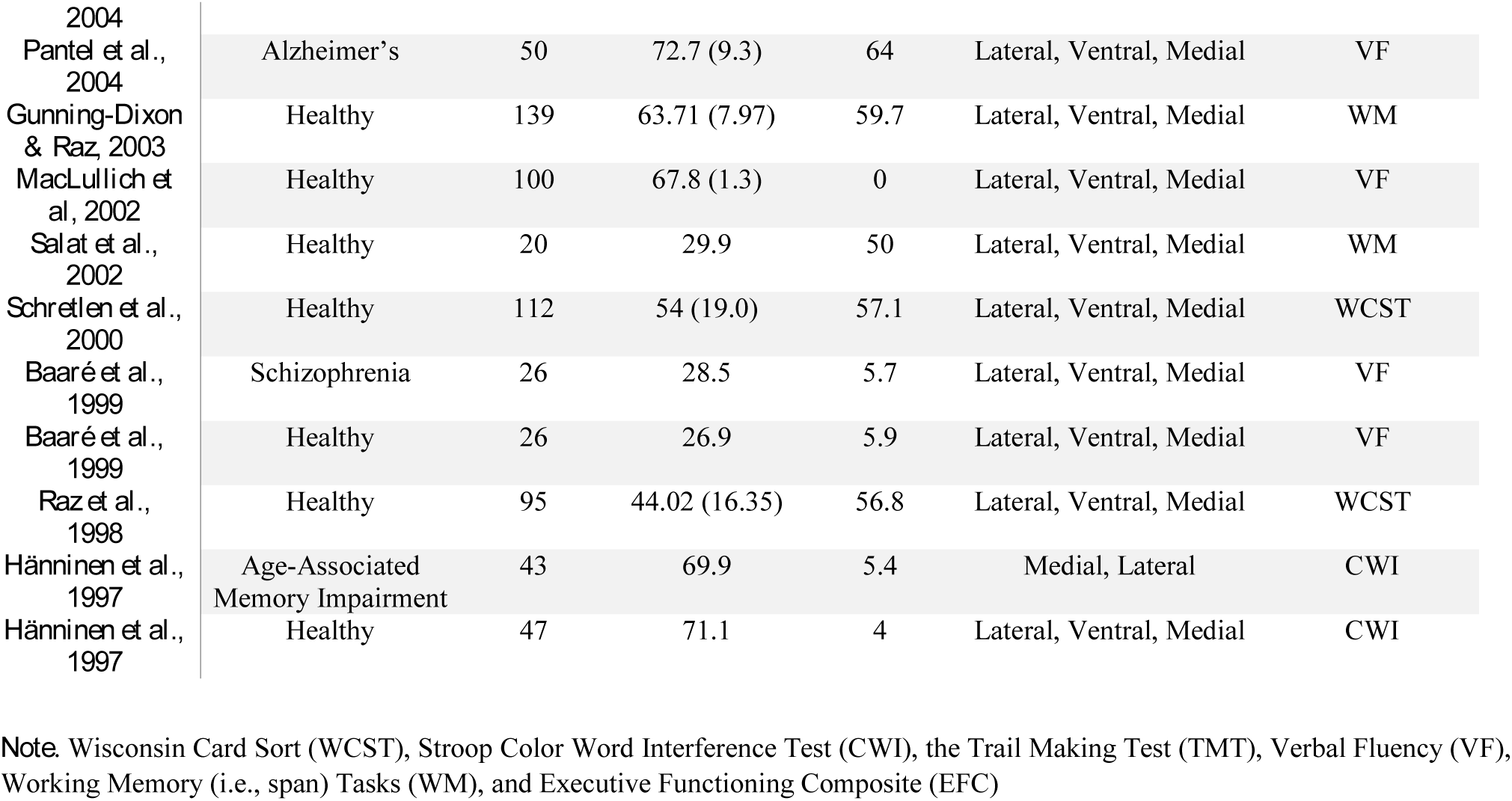
Coded variables for independent samples.

In studies with both healthy controls and neuropsychiatric participants, data from all available groups were utilized, coded for subject type, and separated by group. Effect sizes were calculated for independent samples, for controls and neuropsychiatric patients separately. Due to between-study variability in the operationalization of brain regions, specific ROIs (i.e., orbitofrontal cortex) were not investigated in the present study. The existing level of detail was adequate to support creation of a useful dichotomy: studies were categorized according to whether broad (i.e., diffuse) or specific (i.e., focal) aspects of the prefrontal cortex were operationalized.

### Coding of Variables

For each sample, the following variables were extracted and coded: year, sample type (i.e., healthy or neuropsychiatric), sample size, mean age, standard deviation of age, age range, percentage female, percentage left handed, country of origin, EF task, correlation coefficient, and frontal region. Brain regions were categorized into three non-mutually exclusive groups: medial, lateral, and/or ventral. Brain regions that included volume or thickness from medial, lateral, *and* ventral surfaces were determined to be “broad”. Brain regions that included volume or thickness from *either* medial, lateral, *or* ventral surfaces were determined to be “specific”. This was done to account for between-study variability in the operationalization of regions of interest. EF tasks were coded as one of the following: CWI, VF, WM, TMT, WCST, or an EF Composite score (EFC). Studies with multiple cognitive tasks or multiple brain regions were collapsed under a single effect size, by averaging Fisher’s *z*-scores, for Hypotheses 1 and 4. To compare effect sizes between tasks and brain regions (Hypotheses 2 and 3), these effect sizes were also considered separately. EF measures were reverse coded as needed, so that all positive effect sizes indicated that better performance on EF tasks was associated with larger volume or thickness in frontal brain regions.

#### Data Extraction

Data extraction procedure was designed by senior authors in collaboration with primary coders. Titles were reviewed and excluded by senior author (DG) and reviewed by author AW, with 99.3% agreement. Abstracts were then coded as “excluded”, “included”, and “unsure” by author AW, with senior authors serving as secondary reviewers as needed. Abstracts were coded as “unsure” when there was insufficient information in the abstract to determine if a study should be excluded, and the full article was pulled for review. 248 articles were retained for further examination and data extraction. A primary coder at the undergraduate level was trained by author AW to determine eligibility and extract coded variables. The 248 studies were randomly divided between coder and author AW, and independently coded. To assess interrater reliability, both raters independently coded approximately 10% (*N* = 25) of the 248 remaining studies. After training, the kappa coefficient for eligibility determination was.92. The kappa coefficients and interclass correlation coefficients for extracted variables was 1.0. Author AW reviewed every 10th article to maintain quality control, and reviewed more difficult cases as needed (i.e., determination of statistics eligibility).

### Effect Size Calculation

All correlation coefficients (i.e., *r*) were transformed into *Z*_*r*_ to determine the effect size for each sample. Mean effect size (*M*_*ES*_) was computed for groups by weighting (*w*) each effect size by its sample size (See appendix for formulas). For each *M*_*ES*_, a standard error was calculated and a *z* statistic was generated to determine if the association between executive functioning and brain regions was significant. 95% CIs were calculated using standard errors.

In order to draw more accurate conclusions about the population mean and generalize findings beyond these specific samples, random effects modeling was used. Effect sizes were tested for homogeneity (Hedges’ homogeneity test) to confirm that random effects modelling would be appropriate. Both the healthy samples, χ^2^ (19) = 165.20, *p* <.001, and the neuropsychiatric samples, χ^2^ (20) = 52.47, *p* <.001, were determined to be heterogeneous, which supports the use of random effects modelling. However, because the estimation procedure of random effects modelling can result in bias for samples of interest with less than 10 studies, both fixed and random effects models are reported for the comparison of cognitive tasks, region diffusivity, and specific neuropsychiatric group.

### Random Effects Modelling

For the random effects model (*T_*i*_ = µ+ ξ_*i*_ + ε_*i*_*), τ^2^ (between study variance) was calculated to equal the variance of ξ_*i*_ across the population of studies(18). The between-study variance and subject-sampling variance estimate (*SE*) were used to compute new weights for random-effects analyses. These new weights were used to determine the mean effect sizes (*M*_*ES*_*\*)* for the heterogeneous population.

A *z* statistic was generated to determine if the association between EF and brain regions was significant in the random effects model. Standard errors using these new weights were used to calculate 95% CIs. Mean effect sizes generated from random effects modelling were compared using the observed *z* test statistic and determining if CIs overlapped.

### Fail-Safe *N*

There is a chance in meta-analyses that results are biased by the “file-drawer effect”, where non-significant results are not generally reported. Although this study included non-significant effects as part of these analyses, it is still possible that the selected studies do not effectively represent the true population statistic. A “fail-safe *N*” statistic was computed as an estimate of the number of studies with null results (i.e., calculated effect sizes equal to zero) that would need to be included in analyses to reduce the effect size to a small effect size(19, 20) (*r* = 0.10).

## Results

### Question 1

Larger volumes and thickness were significantly associated with better EF, *z* = 12.53, *p* <.001, in the full sample (*r* =.41, 95% CI =.33 -.48), when effect sizes are collapsed across EF tasks and brain regions (See Figure 2). A fail-safe *N* statistic, determined that an additional 144 studies would be needed to reduce this effect to non-significant levels (*r* =.10).

**Figure 2.**
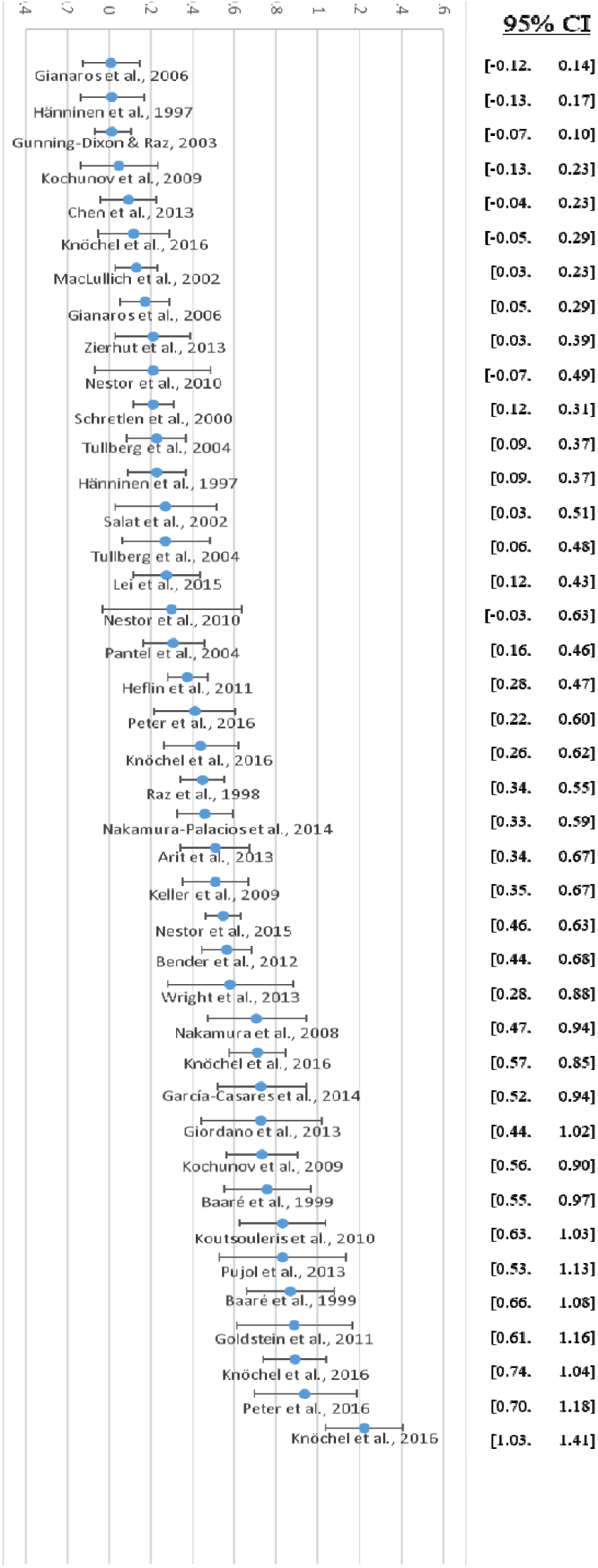
Effect sizes (Z_r_) for associations of cortical volume and executive functions, *z* = 12.53, *p* <.001, in the full sample (*r* =.41, 95% CI =.33 -.48)

### Question 2

#### Cognitive tasks

Effect sizes specific to cognitive tasks varied, with the largest difference between the CWI and EFC (*Z*_*observed*_ = 2.12, *p* <.05). However, there were no significant differences in effect size between specific executive functioning tasks, as evidenced by overlapping CIs. Due to the limited sample studies, both fixed and random effects are reported. Fixed Effects can be found in Table 2 and Random Effects in Table 3.

**Table 2.** Fixed effects for associations between cognitive tests and cortical volume.

|  | *k* | *N* | *Weighted Effect Size* |  |  | 95% CI for *M<sub>ES</sub>* |  |
| --- | --- | --- | --- | --- | --- | --- | --- |
|  |  |  | *M<sub>ES</sub>* | *se* | *p* |  |  |
| Color Word Interference | 5 | 313 | 0.52 | .12 | <.001 | 0.40 | 0.63 |
| Trails | 10 | 525 | 0.48 | .09 | <.001 | 0.39 | 0.57 |
| Wisconsin Card Sort | 5 | 593 | 0.43 | .08 | <.001 | 0.35 | 0.52 |
| Working Memory | 10 | 437 | 0.39 | .10 | <.001 | 0.29 | 0.49 |
| Verbal Fluency | 8 | 351 | 0.37 | .11 | <.001 | 0.26 | 0.48 |
| Composite | 5 | 213 | 0.33 | .14 | <.001 | 0.19 | 0.47 |

*Note.* Overall fixed effect size ( $r = .38$ , 95% CI = .33 - .45)

**Table 3.**
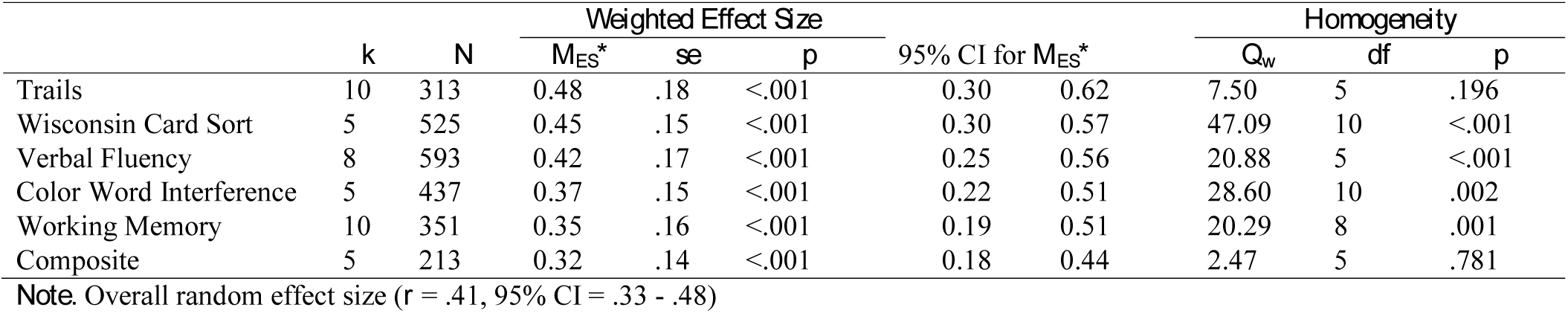
Random effects for associations between cognitive tests and cortical volume.

#### Strength of the association in specific brain regions

Specific brain regions (single brain surfaces, i.e., ROIs that contained either medial, lateral, or ventral areas) had similar effects (*r* =.47, 95% CI [.34-.59]) compared to more diffuse brain regions (three brain surfaces, i.e., ROIs that contained medial, lateral, and ventral areas; *r* =.35, 95% CI [.27-.44]).

### Question 3

#### Healthy and Neuropsychiatric Groups

Separated by sample type, larger volumes and thickness were associated with better executive functioning in both healthy (*r* =.35, 95% CI =.29 -.39) and neuropsychiatric populations (*r* =.47, 95% CI =.40 -.51). The mean effect size was significantly larger for the neuropsychiatric populations (*Z*_*observed*_ = 3.01, *p* <.001), which is consistent with the CIs, which do not overlap. The mean effect sizes for healthy and neuropsychiatric groups can be found in Figure 3.

**Figure 3.**
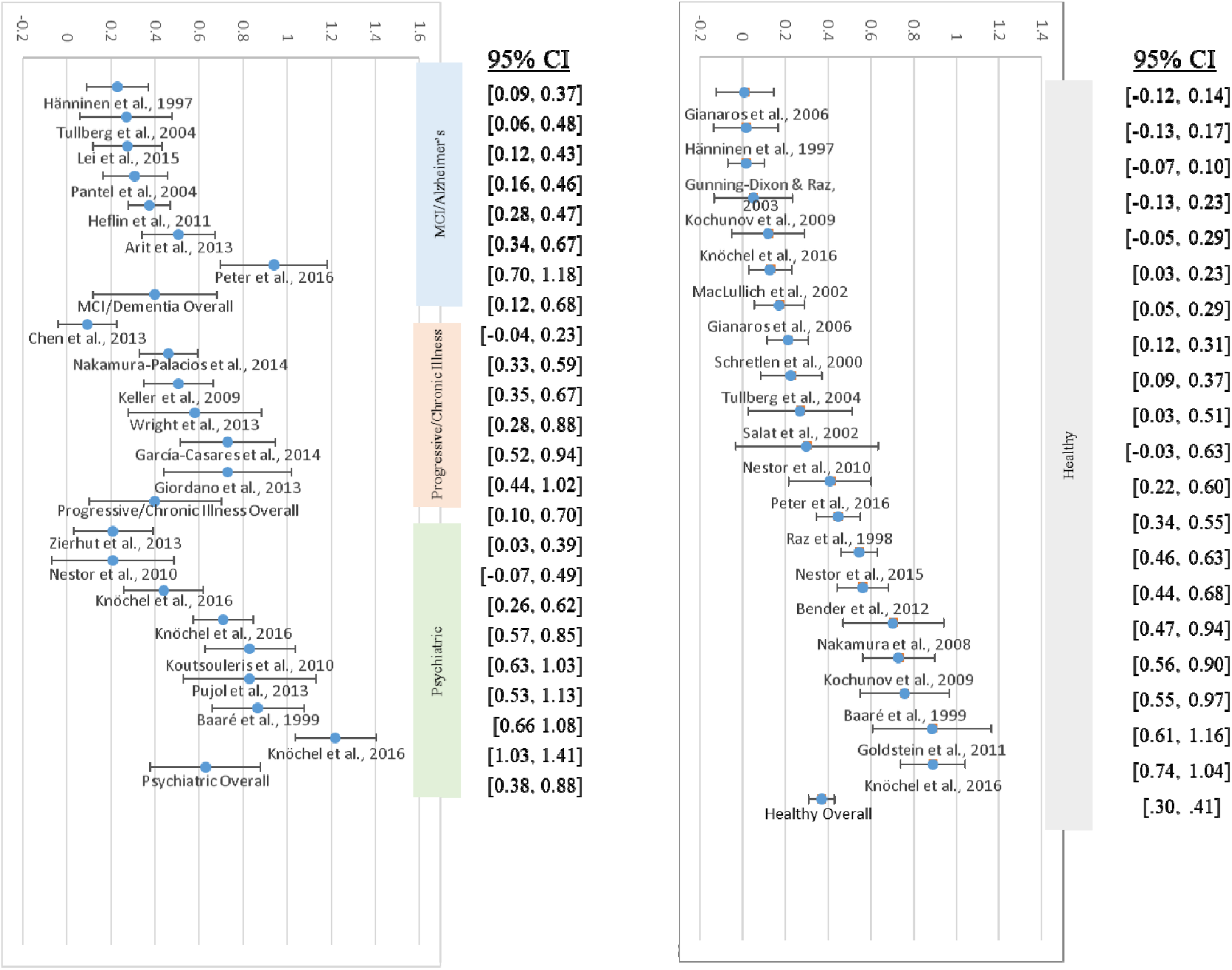
Effect sizes (Z_r_) for associations of cortical volume and executive functions, separated by group.

### Question 4

#### Neuropsychiatric Groups

In order to investigate the variability in the neuropsychiatric group, these were coded as Psychiatric (i.e., Schizophrenia, Bipolar), MCI/Alzheimer’s, or Progressive/Chronic Illness (i.e., progressive supranuclear palsy, temporal lobe epilepsy). Larger volumes and thickness were associated with better EF in Psychiatric (*r* =.56, 95% CI =.36-.71), MCI/Alzheimer’s (*r* =.38, 95% CI =.12-.59), and Progressive/Chronic Illness (*r* =.38, 95% CI=.10-.60) groups. Although the mean effect size for the Psychiatric group trended toward being larger than the MCI/Alzheimer (*Z*_*observed*_ = 2.56, *p* <.05) and Progressive/Chronic Illness groups (*Z*_*observed*_ = 2.03, *p* <.05), the overlapping CIs imply that the effect sizes are not significantly different.

## Discussion

The principal findings of our study were that: (1) there is a significant positive association between EF and cortical size in frontal cortex, and (2) while the magnitude of this effect does not vary as a function of neuropsychological measurement paradigm or specificity of brain region, (3) neuropsychiatric samples have significantly stronger associations between EF and cortical size compared to healthy samples, with volume accounting for 22 and 12%, respectively of EF individual variability.

Our findings regarding the positive association between EF and cortical size confirm findings from a large body of literature examining this relationship. Although the effect size is moderate, the magnitude of the effect is lower than those found in meta-analysis comparing healthy controls and lesion patients(4) (*d* = -.78) and relatively higher than meta-analysis of healthy samples alone(6) (*r* =.15, *d* =.31), both of which were classified as moderate effect sizes. However, these studies generally support a “larger is more powerful” framework(21) to explain relationships between EF and the brain. Our findings regarding this positive relationship between prefrontal cortex structure and EF are also consistent with a meta-analysis of fMRI which found support for a superordinate cognitive control network found in the prefrontal cortex(22), validating structural associations with functional findings. Given the high Fail-safe *N*, it is likely that although the true population effect size is lower than the one estimated in our meta-analysis, it is still of clinically relevant strength. As neuroimaging research continues to move toward “big data” approaches to brain behavior relationships, we will likely see more conservative effect sizes, that may be the target of future meta-analysis. Although this study investigates EF in the context of structural frontal regions, future research should further examine the interaction between functional and structural brain measures as they relate to EF.

The null findings concerning differences between neuropsychological paradigms are not inconsistent with emerging literature support for a robust common EF principle and high heritability of common EF(7). The relationship between volume and any one measure of EF is a function of sensitivity, and these results suggest that these tasks do not vary in their sensitivity to frontal volumes in healthy controls or non-lesion patient samples. Although specific factors of EF have been extracted via factor analysis(23), and meta-analytic review of individual neuropsychological paradigms show specific associations(16, 17), our findings are consistent with lesion studies(4) that show limited comparative differences between task and brain regions as a function of magnitude and effect size (i.e., no one task is more associated with brain volume than another). Although this lack of regionally specific sensitivity does not prove that each paradigm is measuring the same construct, it does suggest that there is a common contribution among paradigms associated with frontal volume.

Of note, the magnitude of the effect size for EFC, while not significantly lower than specific neuropsychological paradigms, did rank last in both the fixed and random effects model. This may, in part, be due to the variability in creation of composite scores. While many of the EFC included in this meta-analysis reported correlations from the overall score of the Frontal Assessment Battery, many contemporary studies utilize a common EF factor derived from factor analysis(24, 25). Aggregate scores, like those found in the Frontal Assessment Battery, aggregate multiple modalities in stimulus presentation and response modality in tasks (i.e., task impurity), as well as some specific executive functioning aspects. Factor analysis extracts those aspects that are common to all tasks, effectively dealing better with task impurity. However, these factor-analytic composites are underrepresented in meta-analyses due to the lack of reported zero-order correlations. The use of composite scores has increased due, in part, to assist in clarity of data interpretation and transdisciplinary communication. As the use of factor-derived composite scores becomes more common place, it is imperative that investigators report their results in ways that facilitate the use of meta-analysis.

Results did not identify either specific or diffuse brain regions as more related to EF. While the random effects model suggests that the magnitude of effects is larger for EF tasks associated with more specific regions, we conclude that the differences are not significant because of large overlap in CIs. Given that there is support in the lesion literature showing that more diffuse damage is associated with larger deficits in EF(9), it would follow that a more diffuse network of regions could also be more strongly associated with EF. It is likely that the effect of the extent of involved tissue is less robust when the pathogenic mechanism is atrophy rather than frank lesion. It is also possible differences between regions are not manifested because of how this study defined the ROIs in order to account for between study variability in cortical regions.

The most salient finding to clinical neuroscientists in this study, is the observed difference in effect size magnitude between healthy and neuropsychiatric populations. This difference is robust enough that despite conservative estimates of confidence intervals as part of random effects modelling, there was no overlap of effects. Because EF tests were originally validated in clinical settings, discrepancies emerge in either the nature or the degree of predictors of performance between populations. These results suggest the relationship between healthy adult performance on neuropsychological testing is less associated with cortical size compared to neuropsychiatric adults. Task performance in healthy controls could be less influenced by properties of the cerebral cortex and more influenced by other factors, such as level of motivation and cognitive reserve, while in neuropsychiatric samples some measured property of the cerebral cortex exerts more important influence. For example, education has been shown in meta-analytic review(26) to significantly predict performance on the TMT (β = −1.31, *se* = 0.44) and VF (β = 0.50, *se* = 0.20) tasks in healthy samples, while it does not significantly predict performance on the Stroop (β = 0.77, *se* = 1.31). Comparisons of cortical integrity between groups, as measured by neuropsychological test performance, should be interpreted with additional caution.

It is also notable that while the neuropsychiatric groups were not significantly different given the presence of overlapping confidence intervals, the effect sizes for the Psychiatric group, which consisted primarily of samples of individuals diagnosed with schizophrenia, trended more robustly than the MCI/Alzheimer’s or Progressive/Chronic Illness groups. The effects for the Progressive/Chronic Illness group may be related to the heterogeneity of this category, which included multiple disease categories. However, longitudinal findings show that treatment resistant patients with schizophrenia have faster rates of age-related cognitive decline than similarly aged patients with Alzheimer’s(27). This may mean that this population is more sensitive to age-related changes in brain structures, particularly in the frontal lobes, which is consistent with differences found here between Psychiatric and MCI/Alzheimer’s groups. These two disease processes have been associated with a number of unique(^28),(29^) and shared(30) genetic factors, which may be contributing to a disparate metabolic processes affecting different brain structures. Although the research regarding specific neural mechanisms of change in schizophrenia is varied, there is meta-analytic support for age-based progressive decreases in grey matter volume(31)and increases in ventricular volume(32). The frontal lobes show the larger decreases relative to global cerebral volume, and the frontal horns of the ventricles show the smallest relative increases(32). This discrepancy suggests that while changes in neural structures related to schizophrenia play a role, comorbid risk factors like substance abuse(33), poor medication compliance(34), or fewer protective factors(35) may also be contributing to reductions in frontal lobe volume, which are more salient for poor outcome patients. Given the transdiagnostic predictive validity of EF, early onset and changes in frontal regions may lead to decreased functional status and its associated risk factors. This may help account for more robust effects in serious mental illness compared to other illness categories, like Alzheimer’s.

There are several limitations to consider regarding the implications of this study. First, for results regarding neuropsychological paradigms, the measurement of effect sizes for the CWI, VF, WCST, and EFC is best represented by the fixed effects given limited sample size (< 10). For samples with fixed effects, conclusions can only be drawn for the studies included, and should be generalized to other samples with caution. Secondly, like all meta-analyses, there is the possibility that our results are influenced by the file drawer effect. Although we identified a moderately large fail-safe N to validate our findings, it is still possible that the generally small sample sizes in imaging studies have contributed to inflated effects. Therefore, it is important to consider group differences as relative to one another, rather than as absolute values of effects.Further research into the variability between neuropsychiatric groups, particularly groups with volumetric deterioration outside of the frontal lobes, will help to determine the specificity of EF tasks as “frontal batteries”. Future use of traditional EF tasks may be oriented more toward validation of biomarkers for future functional decline, and cognitive neuroscience tasks used for localization of function. This may help to clarify specific mechanisms contributing to these group differences, and their relationship to traditional neuropsychological paradigms.

In summary, these findings have clinical implications regarding the interpretability of neuropsychological paradigms as an index of frontal lobe size. Although these tasks have predictive validity for many significant outcomes of interest, it is important to note their limitations in healthy samples, where protective factors such as education may be more predictive than age-related cortical changes(26). This study contributes to meta-analytic findings regarding these brain-behavior relationships and has shown that there are significant changes in effect size magnitude between healthy and neuropsychiatric groups, relative to each other.

## Acknowledgements

The authors would like to acknowledge the following people and organizations for their contributions:

The Suffolk University Psychology Department for their support of doctoral students and David Gansler’s Lab, and the contributions of post-baccheloreate student Mrs. Valeria Vilomar.

## Conflicts of Interest

The authors declare that they have no conflicting interests. This research did not receive any specific grant from funding agencies in the public, commercial, or not-for-profit sectors.

